# The effect of object type on building scene imagery – an MEG study

**DOI:** 10.1101/2020.08.04.236554

**Authors:** Anna M. Monk, Gareth R. Barnes, Eleanor A. Maguire

## Abstract

Previous studies have reported that some objects evoke a sense of local three-dimensional space (space-defining; SD), while others do not (space-ambiguous; SA), despite being imagined or viewed in isolation devoid of a background context. Moreover, people show a strong preference for SD objects when given a choice of objects with which to mentally construct scene imagery. When deconstructing scenes, people retain significantly more SD objects than SA objects. It therefore seems that SD objects might enjoy a privileged role in scene construction. In the current study, we leveraged the high temporal resolution of magnetoencephalography (MEG) to compare the neural responses to SD and SA objects while they were being used to build imagined scene representations, as this has not been examined before using neuroimaging. On each trial, participants gradually built a scene image from three successive auditorily-presented object descriptions and an imagined 3D space. We then examined the neural dynamics associated with the points during scene construction when either SD or SA objects were being imagined. We found that SD objects elicited theta changes relative to SA objects in two brain regions, the right ventromedial prefrontal cortex (vmPFC) and right superior temporal gyrus (STG). Furthermore, using dynamic causal modelling, we observed that the vmPFC drove STG activity. These findings may indicate that SD objects serve to activate schematic and conceptual knowledge in vmPFC and STG upon which scene representations are then built.

## INTRODUCTION

Our lived experience of the world comprises a series of scenes that are perceived between the interruptions imposed by eye blinks and saccades. Indeed, scene mental imagery has been shown to dominate when people engage in critical cognitive functions such as recalling the past, imagining the future and spatial navigation (Clark et al., 2020; Andrews-Hanna et al., 2010; see also Clark et al., 2019). It is not surprising, therefore, that visual scenes have been deployed extensively as stimuli in cognitive neuroscience.

A scene is defined as a naturalistic three-dimensional spatially-coherent representation of the world typically populated by objects and viewed from an egocentric perspective (Dalton et al., 2018; Maguire and Mullally, 2013). Neuroimaging and neuropsychological studies have identified a number of brain areas that seem to be particularly engaged during the viewing and imagination of scenes including the ventromedial prefrontal cortex (vmPFC; Zeidman et al., 2015a; Bertossi et al., 2016; Barry et al., 2019a), the anterior hippocampus (Hassabis et al., 2007a, 2007b; Summerfield et al., 2010; Zeidman et al., 2015a, 2015b; Dalton et al., 2018; reviewed in Zeidman and Maguire, 2016), the posterior parahippocampal cortex (PHC; Epstein and Kanwisher, 1998; reviewed in Epstein, 2008 and Epstein and Baker, 2019), and the retrosplenial cortex (RSC; Park and Chun, 2009; reviewed in Epstein, 2008; Vann et al., 2009; Epstein and Baker, 2019). How are scene representations built, and what specific roles might these brain regions play? While spatial aspects of scenes have been amply investigated and linked to the hippocampus (Byrne et al., 2007; Morgan et al., 2011; Epstein et al., 2017; Epstein and Baker, 2019), the higher-order properties of objects within scenes have received comparatively less attention (Auger et al., 2012; Troiani et al., 2014; Julian et al., 2017; Epstein and Baker, 2019), and yet they could influence how scene representations are constructed by the brain.

One object attribute that seems to play a role in scene construction was reported by Mullally and Maguire (2011; see Kravitz et al., 2011 for related work). They observed that certain objects, when viewed or imagined in isolation, evoked a strong sense of three-dimensional local space surrounding them (space-defining (SD) objects), while others did not (space-ambiguous (SA) objects), and this was associated with engagement of the PHC during functional MRI (fMRI). In other words, SD objects seem to anchor or impose themselves on their surrounding space (e.g., the traffic light in Figure 1A) and in doing so they define the space around them. By contrast, SA objects do not anchor themselves in their surrounding space in the same way (e.g., a worn car tyre in Figure 1A). Participants in the Mullally and Maguire study (2011) were asked to describe the difference between an SD and an SA object, and a typical response included: “SD items conjure up a sense of space whereas SA items float - they go anywhere.” Many participants spontaneously used the word “float” when discussing SA objects, emphasizing the detachment of these objects from an explicit sense of surrounding space. Mullally and Maguire (2011) also found that SD objects tended to be less portable, maintaining a permanent location, although a considerable number of SA objects had reasonably high permanence also (e.g., a large hay bale in Figure 1A), suggesting that permanence, while certainly related to SD, is not the sole basis of the SD effect. Indeed, object permanence has been linked to the RSC rather than the PHC (Auger et al., 2012, 2015; Auger and Maguire, 2013; Troiani et al., 2014). Similarly, Mullally and Maguire (2011) found that object size and contextual associations did not account for the SD effect.

**FIGURE 1.**
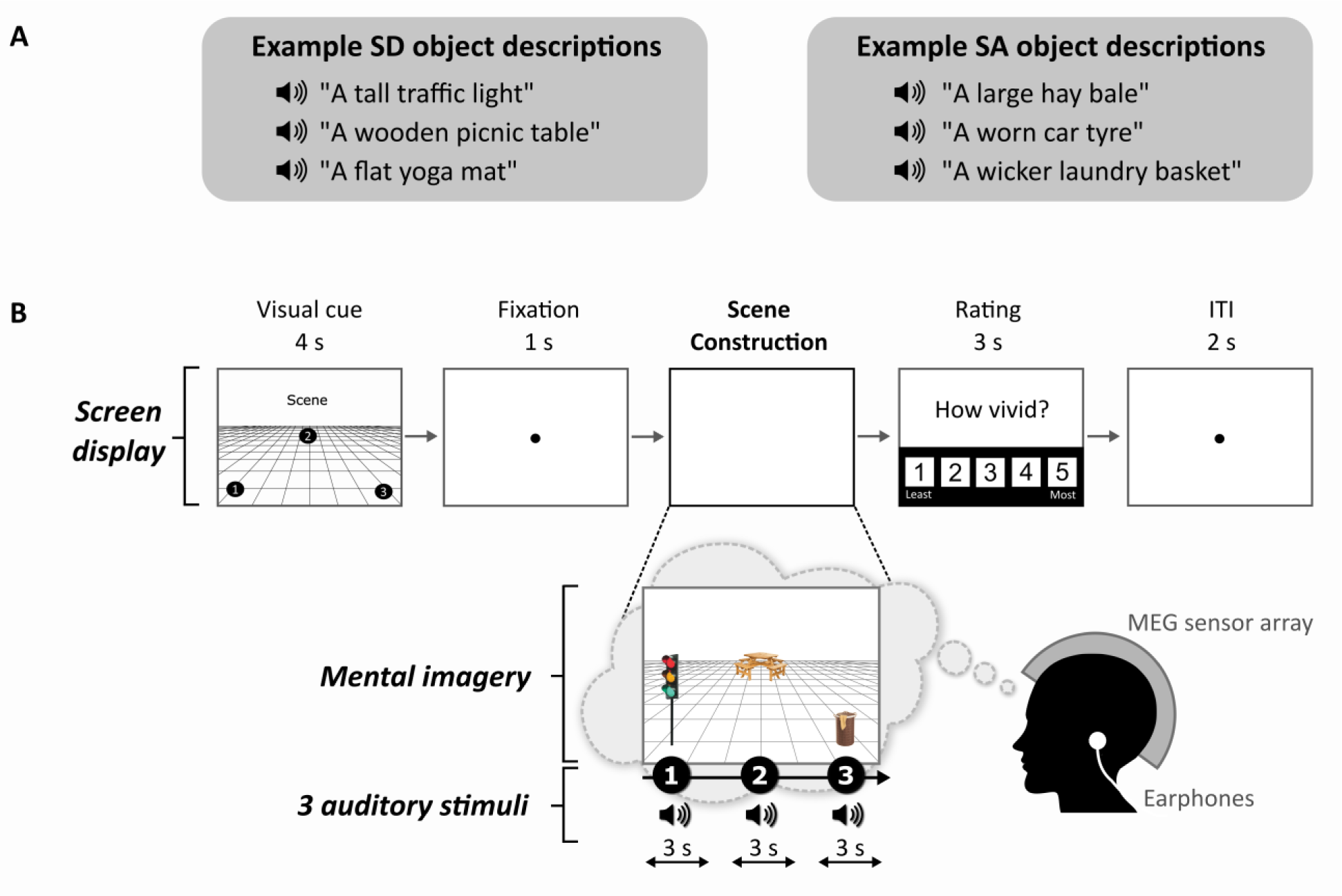
Example stimuli and trial structure. **(A)** Examples of SD and SA object descriptions.**(B)** The structure and timings of an example trial. Note that participants never saw visual objects. During the task the participants imagined the simple scenes while looking at a blank screen.

In a subsequent behavioural study, participants showed a strong preference for SD objects when given a choice of objects with which to mentally construct scenes, even when comparatively larger and more permanent SA objects were available (Mullally and Maguire, 2013). Moreover, when deconstructing scenes, participants retained significantly more SD objects than SA objects. It therefore seems that SD objects might enjoy a privileged role in scene construction.

Mullally and Maguire (2011) examined SD and SA objects in isolation. However, given their apparent influence during scene construction (Mullally and Maguire, 2013), in the current study we compared neural responses to SD and SA objects while they were being used to build imagined scene representations. We adapted a paradigm from Dalton et al. (2018) and Monk et al. (2020) where participants gradually built a scene image from three successive auditorily-presented object descriptions and an imagined 3D space. In order to capture the neural dynamics associated with the points during scene construction when either SD or SA objects were being imagined, we leveraged the high temporal resolution of magnetoencephalography (MEG). In previous MEG studies, changes in vmPFC and anterior hippocampal theta were noted when participants imagined scenes in response to scene-evoking cue words (Barry et al., 2019a, 2019b), and when scene imagery was gradually built (Monk et al., 2020), but the effect, if any, of SD and SA objects on brain responses remains unknown.

Here, we performed a whole brain analysis, and characterized the effective connectivity between any brain regions that emerged from this analysis. While our main interest was in theta, we also examined other frequencies. The obvious prediction, given the previous Mullally and Maguire (2011) fMRI study, was that PHC would be engaged by SD objects. However, because all stimuli were scenes, and the key manipulation of SD and SA objects within scenes was so subtle, we retained an open mind about which brain areas might distinguish between the two object types.

## MATERIALS AND METHODS

### Participants

Twenty-three healthy, right-handed people (13 females; mean age = 25.35 years; standard deviation = 3.69) participated in the experiment. Participants were either monolingual native English speakers, or bilingual native English speakers (i.e., they had English as their first language, but were also but fluent in a second language). They were reimbursed £10 per hour for taking part which was paid at study completion. The study was approved by the University College London Research Ethics Committee (project ID: 1825/005). All participants gave written informed consent in accordance with the Declaration of Helsinki.

### Stimuli

Each stimulus comprised auditorily-presented SD and SA object descriptions (see examples in Figure 1A) and a 3D space. We used the same auditory object descriptions as Dalton et al. (2018). As part of this previous study, the objects were rated as SD or SA. Using these stimuli, we found that SD and SA objects were matched on utterance length (*Z* = 1.643, *p* < 0.1) and number of syllables (*Z* = 1.788, *p* < 0.074). Unsurprisingly, as outlined above, SD objects were rated as more permanent than SA objects (*Z* = 5.431, *p* < 0.001). All objects were rated as highly imageable, obtaining a score of at least 4 on a scale from 1 (not imageable) to 5 (extremely imageable). Objects in each triplet were not contextually related to each other. Participants in the current MEG study were unaware of the SD-SA distinction.

### Task and Procedure

The task, adapted from Dalton et al. (2018) and Monk et al. (2020), involved participants gradually constructing simple scenes in their imagination from a combination of auditorily-presented SD and SA object descriptions and a 3D space. The Cogent2000 toolbox (http://www.vislab.ucl.ac.uk/cogent.php) run in Matlab was used to present stimuli and record responses in the MEG scanner. Auditory stimuli were delivered via MEG-compatible earbuds. Each trial started with a visual cue (4 sec), which displayed the configuration of locations at which objects should be imagined in the upcoming trial (Figure 1B). Four different cue configurations (Figure 2) were randomized across the five scanning blocks. We emphasized the importance of following the cue configurations as precisely as possible. This ensured matched eye movements between the scene and array conditions, consistency across participants, and that objects were imagined as separate and non-overlapping. Participants then fixated on the screen center (1 sec). During the scene construction task (∼9 sec) (Figure 1B), keeping their eyes open whilst looking at a blank screen, participants first imagined a 3D grid covering approximately the bottom two-thirds of the blank screen. Upon hearing each of three auditory descriptions, one at a time, they imagined the objects in the separate, cue-specified positions on the 3D grid. They were instructed to move their eyes to where they were imagining each object on the screen, but also to maintain imagery of previous objects and the grid in their fixed positions. Each construction stage consisted of a ∼2 sec object description and a silent 1 sec gap before the presentation of the next object. An additional 1 sec at the end of scene construction avoided an abrupt end to the task. By the end of a trial, participants had created a mental image of a simple scene composed of a 3D grid and three objects. Vividness of the entire scene was then rated on a scale of 1 (not vivid at all) to 5 (extremely vivid). An inter-trial interval (2 sec) preceded the next trial.

**FIGURE 2.**
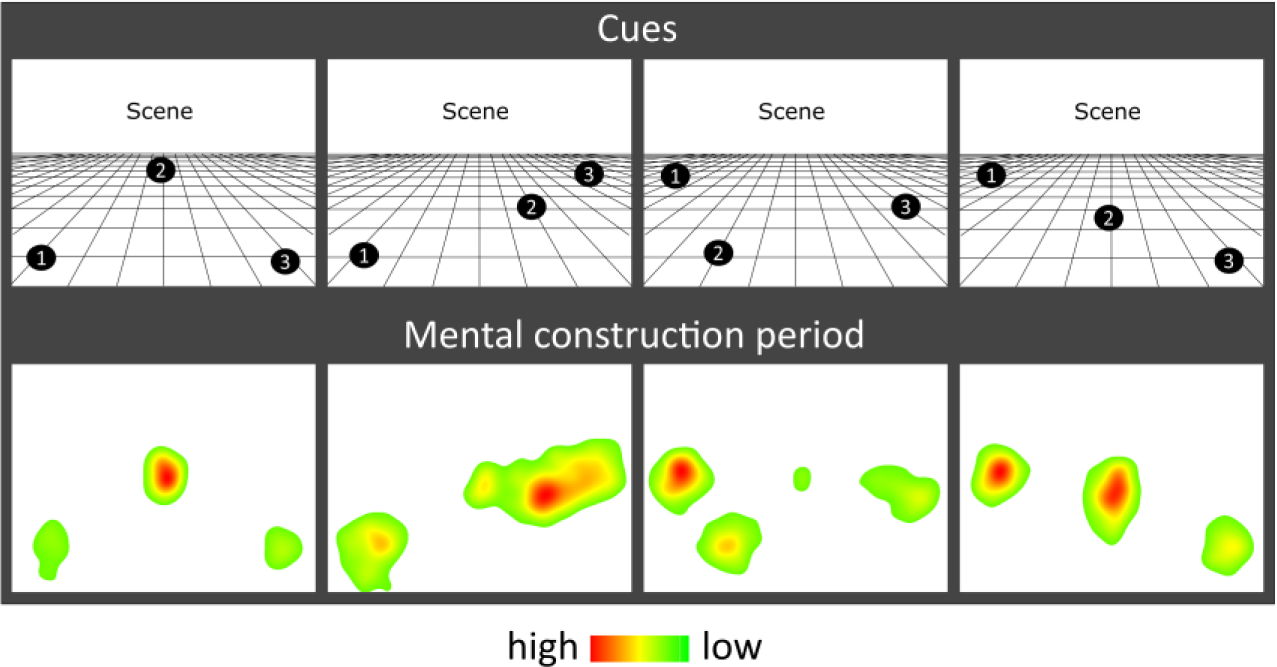
Eye movement results. Heat maps of the fixation count during the 9 sec mental construction period following each cue configuration. Each heat map is an aggregate of fixations on the blank screen across all trials for that cue configuration across all participants with eye tracking data (n=19). Red indicates higher fixation density and green lower fixation density.

Participants imagined a total of 66 scenes (composed of 99 SD, 99 SA objects). Each object description was heard only once. The order of presentation of SD and SA objects within triplets was balanced across scenes with an equal number of SD and SA objects in the first, second and third construction stages (33 in each). A control task (33 trials) involved participants attending to a backward series of auditorily-presented numbers, and was designed to provide relief from the effortful imagination task; it was not subject to analysis. Seven additional catch trials (5 scenes, 2 counting) were also included and pseudorandomly presented across blocks to ensure that participants sustained attention - participants pressed a button upon hearing a repeated object description or number within a triplet.

The visit of each participant took 2.5 hours in total (including training, set up in the MEG scanner and eye tracker calibration, the experimental task (∼50 minutes) and the post-scan memory test).

### Eye Tracking

Eye movements were recorded during the MEG scan using an Eyelink 1000 Plus (SR Research) eye tracking system with a sampling rate of 2000 Hz. The right eye was used for calibration and data acquisition. For some participants the calibration was insufficiently accurate, leaving 19 data sets for the eye tracking analyses.

### Surprise Post-Scan Memory Test

If we found a neural difference between SD and SA objects, it could be argued that this was due to a difference in encoding success. For example, perhaps SD objects were more readily encoded into memory than SA objects, even though there was no explicit memory encoding requirement in this task. By conducting a surprise memory test for SD and SA objects immediately post-scan, we could examine this issue. Participants were presented with a randomized order of all previously heard auditory object descriptions and an additional 33 SD and 33 SA new object description (lures). After hearing each item, they indicated whether or not they had heard the object description during the scan, and their confidence in their decision (1 = low, to 5 = high).

### Behavioral Data Analysis

In-scanner vividness was compared between SD-majority (where 2 out of the 3 objects were SD) and SA-majority scenes (where 2 out of the 3 objects were SA) using a paired-samples t-test. Eye tracking data were analyzed using two-way repeated measures ANOVAs. Memory performance was assessed using the sensitivity index *d’*, based on Signal Detection Theory (Macmillan and Creelman, 1990). We also quantified the bias of participants, to control for any tendency participants might have to give one type of response more than another. This response bias *‘c’* was calculated using the formula: c = - 0.5(Hit Rate + False Alarm Rate). Differences in *d’* and *c* as a function of object type (SD, SA) and construction stage (first, second, third) were each analyzed using a two-way repeated measures ANOVA. Statistical analyses were performed in SPSS25 using a significance threshold of p < 0.05. In cases where Mauchly’s test found sphericity violated, Greenhouse-Geisser adjusted degrees of freedom were applied.

### MEG Data Acquisition and Preprocessing

MEG data were recorded using a 275 channel CTF Omega MEG system with a sampling rate of 1200 Hz. Head position fiducial coils were attached to the three standard fiducial points (nasion, left and right preauricular) to monitor head position continuously throughout acquisition. Recordings were filtered with a 1 Hz high-pass filter, 48-52 Hz stop-band filter, and 98-102 Hz stop-band filter, to remove slow drifts in signals from the MEG sensors and power line interference.

### MEG Data Analysis

All MEG analyses were conducted using SPM12 (www.fil.ion.ucl.ac.uk/spm). Source reconstruction was performed using the DAiSS toolbox (https://github.com/SPM/DAiSS) and visualized using MRIcroGL (https://www.mccauslandcenter.sc.edu/mricrogl).

#### Source Reconstruction

Epochs corresponding to each construction period were defined as 0-3 sec relative to the onset of the SD or SA object description, and concatenated across scanning blocks. Source reconstruction was performed using a linearly constrained minimum variance (LCMV) beamformer (Van Veen et al., 1997). This approach allowed us to estimate power differences between SD and SA objects in source space within selected frequency bands: theta (4-8 Hz), alpha (9-12 Hz), beta (13-30 Hz), and gamma (30-100 Hz).

For each participant, a set of filter weights was built based on data from the SD and SA conditions within each frequency band, and a 0-3 sec time window encapsulating a construction period. Coregistration to MNI space was performed using a 5 mm volumetric grid and was based on nasion, left and right preauricular fiducials. The forward model was computed using a single-shell head model (Nolte, 2003). At the first level, power in a particular frequency band was estimated to create one image per object type (SD or SA) per participant. Images were spatially smoothed using a 12 mm Gaussian kernel and entered into a second-level random effects paired t-test to determine power differences between SD objects and SA objects across the whole brain. An uncorrected threshold of *p* < 0.001 with a cluster extent threshold of >100 voxels was applied to each contrast, in line with numerous other MEG studies (e.g., Barry et al., 2019a; Cushing et al., 2018; Hanslmayr et al., 2011; Kaplan et al., 2012; Kveraga et al., 2011; Lieberman & Cunningham, 2009). This is held to provide a balance between protecting against false positives whilst enabling the detection of subtler signals. The same beamforming protocol was followed when objects were re-categorized as permanent and non-permanent, with the number of permanent and non-permanent objects equalized to 65 in each category.

The contrast of primary interest was the direct comparison between SD and SA objects. As a secondary issue, we were mindful that it was also of interest to know whether an effect represented an increase or a decrease in power from baseline. Including baseline correction in the original beamformer would be the usual way to examine this. However, this is challenging to implement in the case of the current design because an SD or SA object was not necessarily preceded by a clean baseline. On each trial, three object descriptions were heard one after the other. While the first object in a triplet was preceded by a clean 1 sec fixation baseline, objects 2 and 3 had an object imagination stage preceding them. Objects at different construction stages would therefore be corrected against different types of baseline (i.e., object 1 vs preceding fixation, object 2 vs preceding object 1, and object 3 vs preceding object 2). Conducting this analysis could have introduced spurious effects. Consequently, having established that a difference was apparent between SD and SA objects when they were compared directly, we then sought to ascertain the direction of power change. By using a separate beamformer where the 1 sec pre-stimulus fixation period was the only baseline for all objects (whether a SD/SA or first/second/third object), and hence did not overlap with any stimulus period, we were able to establish the direction of power change for SD and SA objects in a straightforward way.

#### Dynamic Causal Modelling (DCM)

Brain areas identified in the whole brain SD versus SA beamformer provided the seed regions for the subsequent effective connectivity analysis, which was conducted using DCM for cross spectral densities (Moran et al., 2009). This approach permitted us to compare different biologically plausible models of how one brain region influences another, as well as mutual entrainment between regions (Friston, 2009; Kahan and Foltynie, 2013). Random-effects Bayesian model selection (BMS) was performed to compare the evidence for each specified model that varied according to which connections were modulated by SD relative to SA objects (Klaas et al., 2009). We determined the winning model to be the one with the greatest exceedance probability. To assess the consistency of the model fit, we also calculated the log Bayes factor for each participant.

## RESULTS

### Behavioral Data

There was no significant difference in the vividness of mental imagery between SD-majority (M = 3.91, standard deviation = 0.69) and SA-majority (M = 3.89, standard deviation = 0.66) scene trials (t_(22)_ = 0.464, p = 0.647). Participants correctly identified on average 97.52% (standard deviation = 0.39) of catch trials, indicating that they attended throughout the experiment.

The effect of object type (SD, SA) and construction stage (first, second, third) on eye-movement fixation count (Fix_Count_) and fixation duration (Fix_Dur_) showed that there were no significant main effects of object type (Fix_Count_: *F*_(1,18)_ = 1.908, *p* = 0.184; Fix_Dur_: *F*_(1,18)_ = 0.086, *p* = 0.772) or construction stage (Fix_Count_: *F*_(2,36)_ = 0.292, *p* = 0.748; Fix_Dur_: *F*_(2,36)_ = 0.535, *p* = 0.590), and no object type×construction stage interaction (Fix_Count_: *F*_(2,36)_ = 0.710, *p* = 0.499; Fix_Dur_: *F*_(2,36)_ = 1.871, *p* = 0.169). Heat maps of the spatial patterns of fixations during the task demonstrated a consistent adherence to cue configuration instructions across participants (Figure 2).

In terms of recognition memory (see Table 1), performance exceeded 80% correct for both SD and SA objects, and for *d’* and *c* there were no significant effects of object type (*d’: F*_(1,22)_ = 0.469, *p* = 0.500; *c: F*_(1,22)_ = 0.012, *p* = 0.915), construction stage (*d’: F*_(2,44)_ = 2.383, *p* = 0.104; *c: F*_(2,44)_ = 0.120, *p* = 0.887), nor were there any interactions (*d’: F*_(2,44)_ = 1.431, *p* = 0.250; *c: F*_(2,44)_ =0.035, *p* = 0.965).

**TABLE 1.**
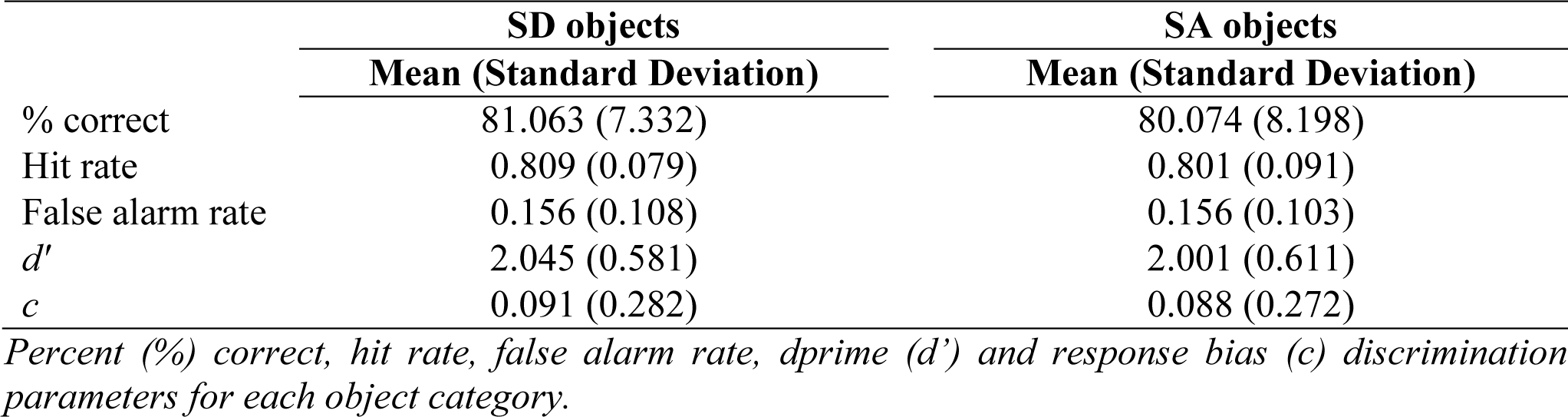
Results of the surprise post-scan object recognition memory test.

### MEG Data

#### Power Changes

A whole brain beamforming analysis revealed significant theta power attenuation for SD compared to SA objects in only two regions: the right vmPFC (peak MNI = 12, 60, −8; *t*-value = 3.66; cluster size = 1960) and right superior temporal gyrus (STG; peak MNI = 66, −6, −12; *t*-value = 3.76; cluster size = 1197) (Figure 3A).

**FIGURE 3.**
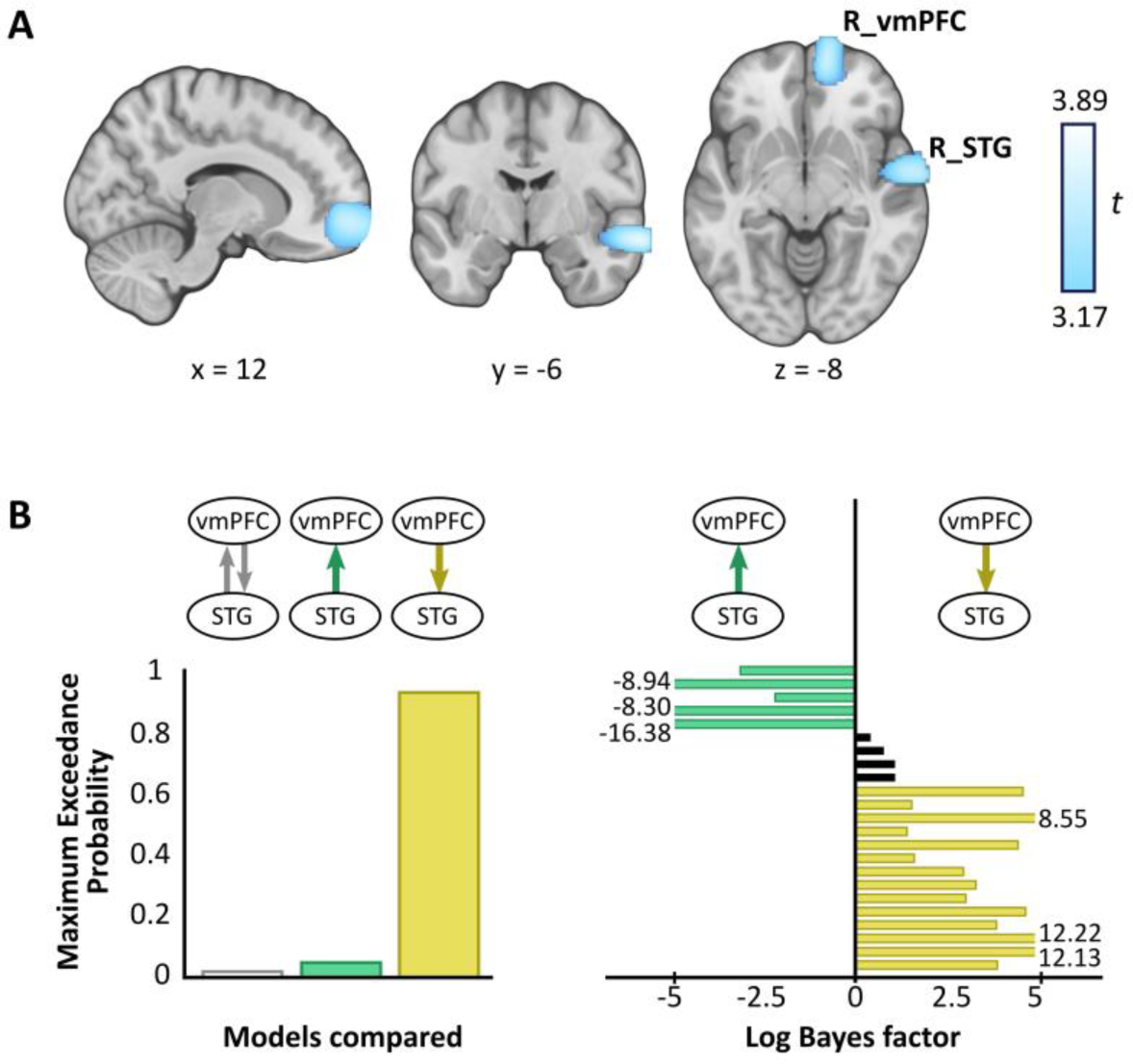
MEG results. **(A)** Source reconstruction of theta (4-8 Hz) power changes during SD relative to SA objects revealed attenuation in the right ventromedial prefrontal cortex (R_vmPFC) and right superior temporal gyrus (R_STG). Images are superimposed on the MNI 152 template and thresholded at uncorrected *p* < 0.001. **(B)** Effective connectivity between R_vmPFC and R_STG was examined using DCM. Three models were compared, with R_vmPFC driving R_STG theta activity during SD compared to SA objects being the model that best explained the data (left panel). Log Bayes factors per participant (right panel) showed positive to strong evidence for this model in most participants. Participants for whom there was no conclusive evidence for either model are represented by black bars. Where log Bayes factors exceeded five, bars are truncated and the exact values are adjacently displayed.

A subsequent contrast between each object type and the baseline revealed that the theta power changes were decreases, echoing numerous previous reports of power attenuation during the construction of scene imagery (e.g., Guderian et al., 2009; Barry et al., 2019a, 2019b) and memory recall (e.g., Solomon et al., 2019; McCormick et al., 2020).

We did not observe any significant differences in theta power between permanent and non-permanent objects across the whole brain.

Analysis of alpha, beta and gamma showed no significant power differences across the whole brain when SD and SA objects were compared.

#### Effective Connectivity

Having established a response to object type in the vmPFC and STG, we next sought to examine the effective connectivity between these regions. We tested three simple hypotheses: (1) vmPFC and STG are mutually entrained, (2) STG drives vmPFC, or (3) vmPFC drives STG. We embodied each hypothesis as a DCM where models differed in which connection could be modulated by SD relative to SA objects. BMS identified the winning model to be vmPFC driving STG during SD more so than SA objects, with an exceedance probability of 91.62% (Figure 3B, left panel). This model was also the most consistent across participants (Figure 3B, right panel).

## DISCUSSION

In this study we focused on an object property, SD-SA, that has been shown to influence how scene imagery is constructed (Mullally and Maguire, 2013). We found that while these object types were being imagined during scene construction, SD objects elicited significant theta changes relative to SA objects in two brain regions, the right vmPFC and right STG. Furthermore, the vmPFC drove STG theta activity.

SD and SA objects were matched in terms of the vividness of mental imagery, eye movements and incidental memory encoding. All objects were incorporated into the same three-object scene structures within which the order of SD or SA object presentation and object locations were carefully controlled. We also examined object permanence, and found that this property did not engage the vmPFC or STG. Our findings are therefore unlikely to be explained by these factors.

Most of the brain regions typically associated with scenes did not respond to SD objects during scene construction. This is likely because scene processing was constant throughout the experiment, and so there was no variation required in the activity of these areas. It is notable that the PHC, which was active during fMRI in response to SD objects when they were viewed or imagined in isolation and devoid of a scene context (Mullally and Maguire, 2011), did not exhibit power changes during scene construction. It may be that examining objects in isolation afforded a “purer” expression of SD whereas, once these objects were included in scene building, higher-order areas then came online to direct their use in constructing scene representations, a possibility that we discuss next.

Considering first the right STG, while this region has been linked to speech processing (e.g., Hullett et al., 2016), the close matching of auditory stimuli and the absence of activity changes in other auditory areas suggests this factor does not account for its responsivity to SD objects. Perhaps more germane is the location of the STG within the anterior temporal lobe, a key neural substrate of semantic and conceptual knowledge that supports object recognition (Peelen and Caramazza, 2012; Chiou and Lambon Ralph, 2016). Patients with semantic dementia, caused by atrophy to the anterior temporal lobe, lose conceptual but not perceptual knowledge about common objects (Campo et al., 2013; Guo et al., 2013).

This could mean that SD objects provide conceptual information that is registered by the STG. Why might this be relevant to scene construction? Prior expectations have a striking top-down modulatory influence on our perception of the world, enabling us to process complex surroundings in an efficient manner (Summerfield and Egner, 2009), and resolve ambiguity (Chiou and Lambon Ralph, 2016). Without this knowledge, we are unable to understand how and where an object should be used (Peelen and Caramazza, 2012). Therefore, objects are an important source of information about the category of scene being imagined (or viewed), facilitating a rapid, efficient interpretation of the scene ‘gist’ without the need to process every component of a scene (Oliva and Torralba, 2006; Summerfield and Egner, 2009; Clarke and Tyler, 2015; Trapp and Bar, 2015). For example, if we see a park bench this might indicate the scene is from a park. Although in the current study the scenes were deliberately composed of semantically unrelated objects, this may not have impeded the STG in nevertheless registering SD objects more so than SA objects because SD objects would normally offer useful conceptual information to help anchor a scene.

The operation of the right STG might be facilitated by the right vmPFC. Converging evidence across multiple studies has shown that the part of the vmPFC that was active in response to SD objects plays a role in the abstraction of key features across multiple episodes (Roy et al., 2012). These contribute to the formation of schemas, which are internal models of the world representing elements that likely exist in a prototypical scene, based on previous exposure to such scenes (Tse et al., 2007; van Kesteren et al., 2013; Gilboa and Marlatte, 2017). For instance, a park typically contains benches, trees and flowers. SD objects may be particularly useful in building scene schema, and hence the response to them by the vmPFC.

Patients with damage to the vmPFC exhibit deficits that suggest aberrant schema re-activation (Ciaramelli et al., 2006; Gilboa et al., 2006; Ghosh et al., 2014), and this has led to the proposal that vmPFC may activate relevant schema to orchestrate the mental construction of scenes performed elsewhere – for example, in the hippocampus (McCormick et al., 2018; Ciaramelli et al., 2019; Monk et al., 2020). Our DCM findings extend this work by showing that the right vmPFC also exerts influence over the right STG, indicating it may be engaging in top-down modulation of conceptual object processing by the STG, specifically during the processing of SD objects. Our results may therefore indicate that SD objects help to define a scene by priming relevant schemas in the vmPFC which then guide conceptual processing in areas such as the STG.

There is another possible explanation for our findings. In the current study, the scenes were deliberately composed of semantically unrelated objects, and this could have introduced ambiguity about a scene’s identity. vmPFC and STG engagement may therefore be evidence of additional neural processing that was required to resolve incongruences inherent to acontextual scenes (Chiou and Lambon Ralph, 2016; Brandman and Peelen, 2017; Epstein and Baker, 2019), perhaps by drawing upon existing schemas in the pursuit of an appropriate scene template. Indeed, connectivity between medial prefrontal and medial temporal cortex has been shown to increase when novel information that was less congruent with pre-existing schematic representations was processed (van Kesteren et al., 2010; Chiou and Lambon Ralph, 2016). The acontextual nature of the scenes, and the effortful nature of the scene construction task, may also have precluded observation of a schema-related memory advantage for SD objects (see more on schema and memory in McCormick et al., 2018; Gilboa and Marlatte, 2017; Tse et al., 2007). It should be noted, however, that our study was not designed to investigate schema, and consequently these possible interpretations remain speculative. Future studies will be needed to further elucidate the SD-SA difference revealed here, perhaps by comparing semantically related and unrelated objects during scene construction, and by adapting the current paradigm to test patients with vmPFC or STG damage. Another notable feature of our findings was the right hemisphere location of the responses. This may be related to the visual nature of the imagined scenes. This too could be probed further in future studies by comparing patients with left and right-sided lesions.

In conclusion, this study revealed the neural dynamics associated with a specific object property during scene construction, and we suggest that SD objects in particular may serve to activate schematic and conceptual knowledge in vmPFC and STG upon which scene representations are then built.

## DATA AVAILABILITY STATEMENT

The raw data supporting the conclusions of this manuscript, and the test materials, will be made available by the authors to any qualified researcher upon request. Requests can be sent to.

## ETHICS STATEMENT

The study was approved by the University College London Research Ethics Committee (project ID: 1825/005). All participants gave written informed consent in accordance with the Declaration of Helsinki.

## AUTHOR CONTRIBUTIONS

AMM and EAM designed the study. AMM collected and analyzed the data with input from EAM and GRB. AMM and EAM wrote the manuscript. All authors contributed to manuscript revision, read and approved the submitted version.

## FUNDING

This work was supported by a Wellcome Principal Research Fellowship to EAM (210567/Z/18/Z) and the Centre by a Centre Award from the Wellcome Trust (203147/Z/16/Z).

## ACKNOWLEDGEMENTS

Thanks to Marshall Dalton for providing the task stimuli, Peter Zeidman for his DCM advice, and Daniel Bates, David Bradbury and Eric Featherstone for technical support.

## CONFLICTS OF INTEREST

The authors declare that the research was conducted in the absence of any commercial or financial relationships that could be construed as a potential conflict of interest.

## Notes

### Competing Interest Statement

The authors have declared no competing interest.

